# Hematopoietic stem cells self-renew symmetrically or gradually proceed to differentiation

**DOI:** 10.1101/2020.08.06.239186

**Authors:** Melania Barile, Katrin Busch, Ann-Kathrin Fanti, Alessandro Greco, Xi Wang, Hideyuki Oguro, Qin Zhang, Sean J. Morrison, Hans-Reimer Rodewald, Thomas Höfer

**Author notes:** These authors contributed equally (Melania Barile, Katrin Busch, Ann-Kathrin Fanti).

## Abstract

It is not known whether hematopoietic stem cells (HSCs) undergo symmetric or asymmetric cell divisions in the unperturbed bone marrow. Here, we integrate data from HSC fate mapping and cell-cycle-dependent labeling through mathematical inference and thus gain insight into how HSCs coordinate self-renewal with differentiation. We find that most HSC divisions in adult mice are symmetric self-renewing, replacing HSCs lost by direct differentiation and death, and slowly expanding the HSC population. This expansion maintains constant HSC output to multipotent progenitors (MPPs), despite declining HSC differentiation rate with age. We identify a linear hierarchy of differentiation states between tip HSCs and MPPs, where *Tie2*-driven HSC fate mapping fully covers the progression of the differentiating cells. A turning point from self-renewal to accelerated cell differentiation occurs between early-stage and late-stage MPPs, just before lineage differentiation becomes manifest in single-cell transcriptomes. This stem cell hierarchy precedes lineage differentiation and may limit mutation accumulation in the hematopoietic system.

## INTRODUCTION

Tissue stem cell populations maintain themselves while producing differentiated output. This balance was often thought to be realized by stem cells dividing asymmetrically into a stem and a progenitor cell (Osgood, 1957; Inaba and Yamashita, 2012). However, stem cell populations are sustained in the face of cell death or even expand, arguing for a role of symmetric self-renewing divisions under physiological conditions (Morrison and Kimble, 2006). There is now considerable evidence for symmetric stem cell self-renewal in solid tissues (Simons and Clevers, 2011; Klein and Simons, 2011). By contrast, how stem cells in hematopoiesis – one of the most productive tissue renewal processes – combine maintenance and productivity *in vivo* remains poorly understood (Höfer and Rodewald, 2016; Laurenti and Gottgens, 2018). Early theoretical work hypothesized “division of labor” between symmetrically self-renewing hematopoietic stem cells (HSCs) and HSCs differentiating into progenitor cells (Lajtha et al., 1962). Recently, observation of asymmetric protein segregation *in vitro* indicated that a minority of HSC divisions (~20%) are asymmetric while the large remainder are symmetric self-renewing (Loeffler et al., 2019). Whether these findings extrapolate to HSC behavior in the intact bone marrow is not clear.

Understanding the mechanisms that regulate stem cell maintenance and output requires methods to quantify their behavior *in vivo*, without disrupting stem cell interactions with their niches (Morrison and Spradling, 2008; Mende et al., 2019). Cell-cycle-dependent labeling studies reported that HSCs divide infrequently (between 2 and 7% of HSCs per day) and indicate that they are not a uniform population in this respect (Kiel et al., 2007; Wilson et al., 2008; Foudi et al., 2009; Oguro et al., 2013; Akinduro et al., 2018). The least proliferative (“proliferation-label retaining”) HSCs have been termed dormant (Wilson et al., 2008; Bernitz et al., 2016; Cabezas-Wallscheid et al., 2017), but careful analysis of label dilution has shown that even these cells cycle at low rate (Morcos et al., 2020). Cell-cycle-dependent labeling approaches could not disentangle asymmetric and symmetric self-renewal of HSCs nor assess HSC differentiation.

Recently, Cre-lox-based approaches for the genetically heritable labeling of HSCs in their niches *in vivo* have become available, allowing the fate mapping of HSC populations during physiological hematopoiesis (Gazit et al., 2014; Busch et al., 2015; Sawai et al., 2016; Chapple et al., 2018). Combining fate mapping after selective HSC labeling with mathematical inference, we previously reported that about 1% of HSCs per day give rise to multipotent progenitor cells (MPPs, named also short-term repopulating HSCs, or ST-HSCs) (Busch et al., 2015). While this low frequency of HSC differentiation has been controversially discussed based on fate mapping experiments with less specific initial labeling of both HSCs and progenitors (Sawai et al., 2016; Säwen et al., 2018; Chapple et al., 2018), a variety of experimental studies concluded that, indeed, adult hematopoiesis functions for extended time periods (at least half a year) independent of HSCs (Sun et al., 2014; Schoedel et al., 2016; Sheikh et al., 2016; Pei et al., 2017; Rodriguez-Fraticelli et al., 2018).

Hence, HSC proliferation and differentiation *in vivo* have so far been studied separately using either cell-cycle-dependent (“diluting”) labeling and HSC fate mapping with inducible, genetically heritable labels. “Microscopic” stem cell decisions, such as asymmetric division versus symmetric self-renewal, have remained beyond the resolution of these population-level approaches. Here, we reason that insight into these fundamental stem cell decisions *in vivo* will be gained by joint analysis of the two kinds of labeling data. Principally, this should be the case as both proliferation and differentiation of hematopoietic stem and progenitor cells (HSPCs) control their numbers, and all three features can be measured, via cell-cycle-dependent labeling (proliferation), via fate mapping (differentiation), or directly (cell numbers). We show that mathematical balance equations that link to cell numbers to fundamental stem cell processes enables statistical inference of the rates of these processes from experimental data. Our inference framework accommodates temporal changes in the rates of the stem cell decisions and, therefore, can quantify HSC behavior during aging. We find that HSC divisions in adult hematopoiesis are predominantly symmetric self-renewing, which is key to maintaining a constant output of MPPs throughout life in the face of a declining HSC differentiation rate. Down-stream of HSCs, extensive self-renewal *in vivo* persists well into the MPP stage. Juxtaposing data from fate mapping and single-cell RNA-seq, we find that self-renewal is progressively lost within late-stage MPPs, just before lineage differentiation becomes discernable in the transcriptomes of individual cells.

## RESULTS

### Opposing gradients of heritable and cell-cycle-dependent labels in hematopoietic stem and progenitor cells

Hematopoietic stem cells have phenotypically been defined as Lin^−^Kit^+^Sca1^+^ (LSK) CD150^+^ CD48^−^ cells, which give rise to MPPs (LSK CD150^−^CD48^−^) and hematopoietic progenitor cells (HPCs, LSK CD48^+^). To gain finer resolution, we here follow Oguro et al. (2013) who introduced additional surface markers to subdivide HSCs (into HSC1 and HSC2) and MPPs (into MPP1-3), with multilineage reconstitution potential after transplantation declining gradually from HSC1 to MPP3 and being lost in HPCs (Figure 1A). We introduced heritable YFP expression by treating our previously reported *Cre* knockin mice, *Tie2*^*MeriCreMer*^*Rosa26*^*YFP*^ (expressing codon-improved Cre flanked by modified estrogen receptor domains from the *Tie2* locus; Busch et al., 2015) with tamoxifen (Figure 1B). This treatment labeled initially HSC1 and HSC2, with preferential labeling of HSC1, but no MPPs or HPC1 (Figure 1C, 8 days post tamoxifen). Hence HSC labeling was specific and focused on the presumed most primitive stem cells, HSC1, which contrasts with previous fate mapping studies where downstream progenitors were also initially labeled by other Cre drivers (Sawai et al., 2016; Chapple et al., 2018; Säwen et al., 2018).

**Figure 1.**
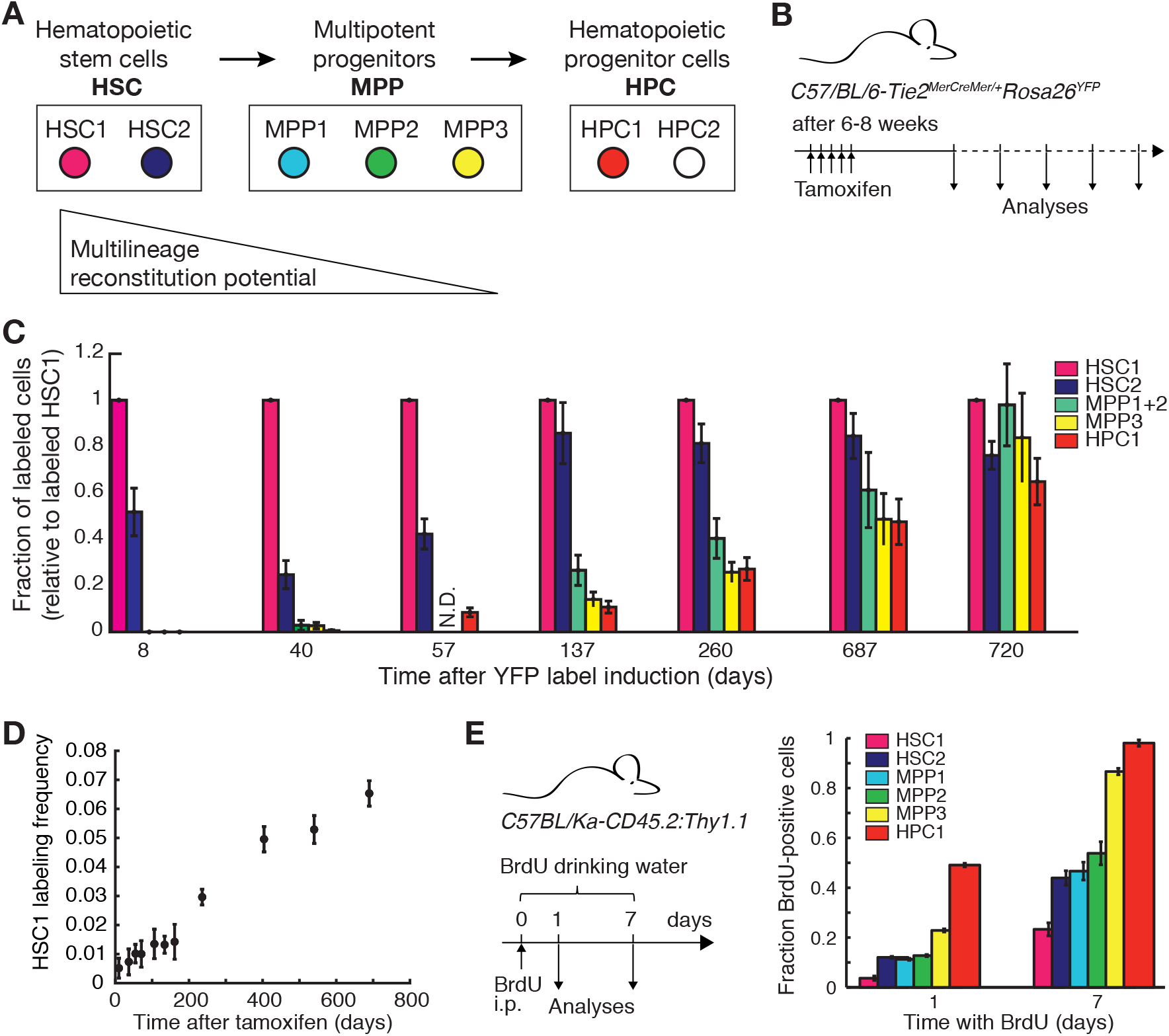
HSC fate mapping, and cell-cycle-dependent labeling of hematopoietic stem and progenitor cells. (A) Assayed hematopoietic stem and progenitor cell populations. HSCs (LSK CD150^+^CD48^−^) and MPP (LSK CD150^−^CD48^−^) were subdivided by the additional SLAM markers CD229 and CD244 as follows: HSC1 (CD229^−^CD244^−^) and HSC2 (CD229^+^CD244^−^); MPP1 (CD229^−^CD244^−^), MPP2 (CD229^+^CD244^−^) and MPP3 (CD229^+^CD244^+^). HPCs are LSK CD48^+^, the vast majority of which are HPC1 (CD150^−^CD48^+^). The reconstitution potential of these cells after transplantation into irradiated recipient mice declines gradually, as reported by Oguro et al. (2013). (B) Fate mapping of HSCs using *C57/BL/6*-*Tie2*^*MCM*^*Rosa26*^*YFP*^ mice. Cre was activated in 6-8 week old mice by tamoxifen injection to switch on heritable YFP expression from the *Rosa26* locus. Mice were sacrificed and cell populations analysed for frequency of YFP^+^ cells at different time points. (C) Propagation of fate-mapping HSC label into hematopoietic stem and progenitors cells. Frequencies of YFP-labeled cells in the bone marrow of one hindleg were measured at the indicated times (*N* = 4-35 mice per time point; error bars indicate standard error of the mean, SEM). For robustness, the relatively rare MPP1 were combined with MPP2. (D) Labeling frequency in HSC1 increases over time, indicating that a subpopulation of tip-HSCs was preferentially labeled. (E) Cell-cycle-dependent labeling of hematopoietic stem and progenitor cells by continuous incorporation of BrdU, administered by intraperitoneal injection into *C57/BL/Ka-CD45.2:Thy1.1* mice and additionally supplied in the drinking water (data redrawn from Oguro et al., 2013; *N* = 5 per time point; error bars, SEM

To examine to what extent HSC1 or HSC2 serve as the source of hematopoiesis under physiological conditions, we performed fate mapping experiments, chasing YFP (fate-mapping) label for up to two years after its induction in young adult mice. The fraction of YFP-positive cells slowly and continuously increased in HSC2 and all progenitor populations relative to HSC1 (Figure 1C). Labeling frequency of HSC2 equilibrated after 4.5 months (Figure 1C, 137 days) with HSC1 while labeling in all MPP subpopulations equilibrated with HSC1 after two years (Figure 1C, 720 days) within measurement error. In absolute terms, labeling frequency of HSC1 increased, from an average of less than 1% immediately after tamoxifen application to nearly 7% two years later (Figure 1D). While previous analysis of HSC up to 200 days showed relatively constant labeling (Busch et al., 2015), the long-term increase in HSC1 seen here became particularly noticeable in aging mice. Taken together, these findings show that HSC1 serve as a persistent source of fate-mapping label, and hence differentiating progenitor cells, throughout the mouse’s life.

Only a fraction of HSC1 was initially labeled, which may simply be due to Cre randomly switching on YFP expression in some HSC1 cells but not others (Pei et al., 2017). However, there could also be a functionally distinct HSC1 subpopulation that is not reached by *Tie2*^*MeriCreMer*^ (*Tie2*^*MCM*^ from hereon) activity, and hence our fate mapping experiments would report only on a subset of HSC1⟶ MPP differentiation pathways. To address this question, we examined the mathematical laws of fate mapping and found that, under general (steady state or non-steady state) conditions, a product population with a unique labeled stem cell source will approach the labeling frequency of this source over time, whereas an additional unlabeled source would keep product label forever below source label (Box 1). As we observe complete label equilibration between all MPP subpopulations and HSC1 (Figure 1C), our fate mapping experiments fully cover the underlying differentiation process, suggest that there is no significant subset of HSCs in which *Tie2*^*MCM*^ does not recombine. We emphasize that this conclusion is independent of the numerical value of the stem cell labeling frequency, which appears to have been unclear in previous work (Sawai et al., 2016).

These conclusions rest on the observation that *Tie2*^*MCM*^ causes persistent labeling of HSC1 and hence the labeled cells are true stem cells. Given that *Tie2*^*MCM*^ activity is restricted to the initial time of tamoxifen application and subsequently ceases (Figure 1B; Busch et al., 2015; Pei et al., 2017), the long-term increase in HSC1 labeling (Figure 1D) suggests that there exists a tip stem cell population with at least ~ 7% labeling frequency that is enriched in HSC1 compared to HSC2. This finding is consistent with a previous study showing that high *Tie2* expression marks tip stem cells with a *Tie2* expression reporter (Ito et al., 2016); however, available anti-Tie2 antibodies did not allow reliable detection by FACS (not shown). The observed label increase in HSC1 suggests that tip HSCs expand with age relative to the remainder of the HSC1 compartment, which we will examine further below.

We found that fate mapping also provides information on precursor-product relations down-stream of the initially labeled stem cells (Box 1). The labeling frequency in a precursor will always be larger than in its product(s), before both eventually equilibrate with the stem cells. This inequality holds even when precursor or product comprise a series of subpopulations that are not recognised by the phenotypic markers used (i.e., the phenotypically defined populations are “heterogeneous”). Hence, the observation that the assayed HSPCs retain the same rank order with respect to their labeling frequencies at all intermediate time points (Figure 1C, 40 days to 687 days) is consistent with the linear differentiation pathway HSC1 ⟶ HSC2 ⟶ MPP1+2 ⟶ MPP3 ⟶ HPC1.

Finally, to assay proliferation of HSPCs, Oguro et al. (2013) used low concentrations of BrdU, so as not to stimulate HSC proliferation (Kiel et al., 2009) (data replotted in Figure 1E). The observed rank order with respect to BrdU incorporation was the mirror image of the gradient in heritable labeling (cf. Figure 1C). These data suggest that progressive differentiation of cells from HSCs towards downstream progenitor stages is accompanied by an increase in cell proliferation.

### Inference of symmetric and asymmetric divisions, direct differentiation and loss of stem cells *in vivo*

We asked which information about HSC behavior can be inferred from the combined fate mapping and BrdU pulse labeling data (Figure 1C and 1E). The number of stem cells results from the balance of five basic processes: symmetric self-renewal, asymmetric cell division, symmetric division giving rise to two differentiating progenitors, direct differentiation to progenitors and cell death, which may not all occur (Figure 2A). The frequencies of stem cells involved in these possible processes per time unit are quantified by rates: *σ* (symmetric self-renewal), *γ* (asymmetric division), *ρ* (symmetric differentiating division), *μ* (direct differentiation) and *δ* (cell death) (Figure 2A), which cannot be observed directly in the intact bone marrow and thus need inference. Propagation of heritable fate-mapping label, and acquisition of a cell-cycle-dependent label (e.g., a thymidine analog) differentially depend on these rates (green and red stars, respectively, in Figure 2A), and hence the combination of the two kinds of data should yield more information than each data type alone. Indeed, by deriving the mathematical equations describing these experimental data (Methods), we found that the following four aggregate rate parameters of the fundamental rates can be inferred (Figure 2B): First, the *total differentiation rate* of stem cells

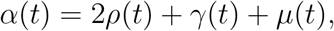

measures how many downstream progenitors are produced per time unit and stem cell. Second, the *net loss rate*

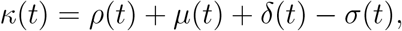

determines whether the stem cell population is maintained (*κ* = 0), shrinks (*κ* > 0), or expands (*κ* < 0, “net gain”) over time. Third, the *total rate of cell divisions*

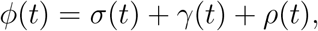

quantifies how often a stem cell divides, regardless of the types of daughter cells. Finally, the *self-renewal rate*

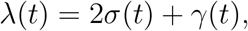

measures how many stem cells are produced by symmetric and asymmetric divisions combined. Of note, the model is suitable for probing changes of HSC proliferation or differentiation over time and does not require stem cell numbers to be constant.

**Figure 2.**
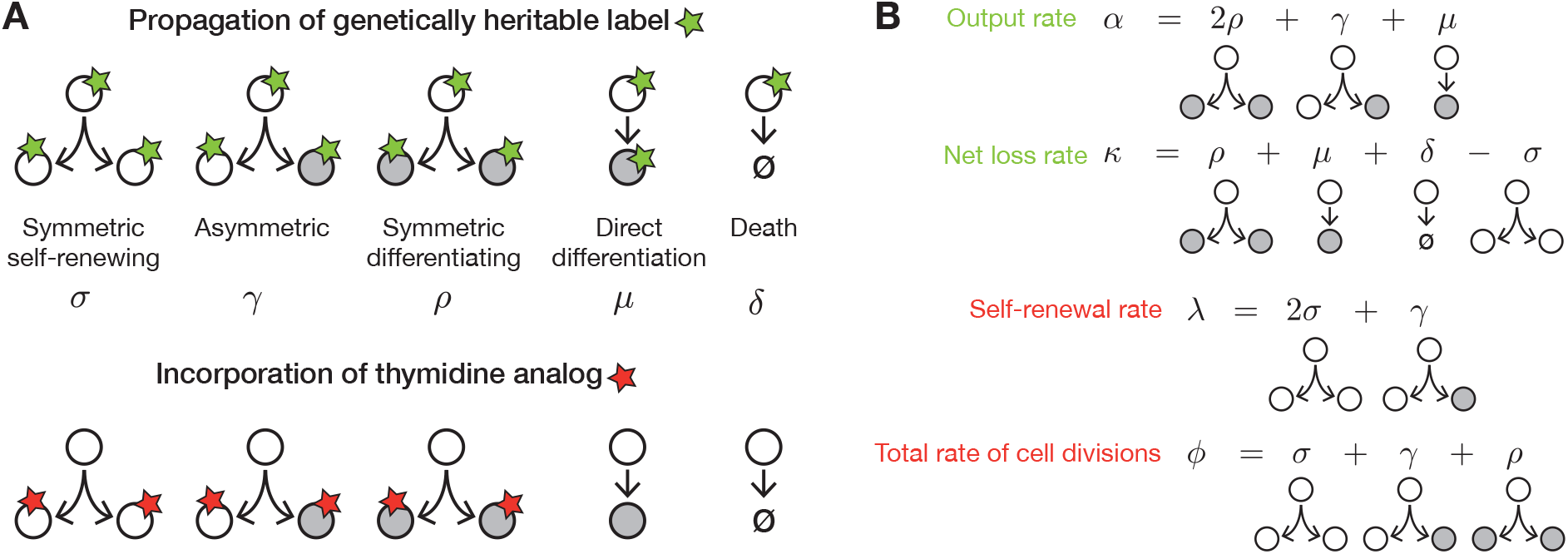
Stem cell decisions quantified by combining fate mapping and cell-cycle-dependent labeling. (A) Basic stem cell decisions (with rates indicated by Greek symbols), with propagation of fate mapping label (green starts, upper row) and incorporation of thymidine analog (red stars, lower row). (B) Aggregate rate parameters obtained by combining fate mapping and thymidine analog pulse labeling. Parameters primarily obtained from fate mapping (thymidine pulse labeling) shown in green (red).

The four aggregate parameters *α*, *κ*, *ϕ* and *λ* are independent (i.e., no parameter can be replaced by a combination of the remaining three). However, they can be combined to yield additional parameters, including the rate of cell death (*δ* = *κ*+*ϕ*−*α*), e.g., by apoptosis (Domen et al., 2000), and the rate of direct differentiation (*μ* = *α* + *λ* − 2*ϕ*). Moreover, symmetric differentiating HSC divisions have not been reported (Loeffler et al., 2019); setting their rate *ρ* to zero one also obtains the rates of symmetric self-renewing and asymmetric HSC divisions (as *σ* = *λ* − *ϕ* and *γ* = 2*ϕ* − *λ*, respectively). Experimental data on stem cell fate mapping will reveal *α*(*t*) and *κ*(*t*) and data on thymidine analog pulse labeling will yield *ϕ*(*t*) and *λ*(*t*) (Figure 2B, Methods) and thus the combined data will provide insight into the rates of fundamental stem cell decisions *in vivo*.

### Hematopoietic stem cells self-renew symmetrically in the bone marrow

We applied this inference framework to the long-standing question whether HSCs self-renew symmetrically or asymmetrically in the bone marrow. To this end, we used a large data set on label propagation from HSCs to MPPs in *Tie2*^*MCM*^*Rosa26*^*YFP*^ mice, labeled in young adulthood and traced up to 8 months of age. Over this time period, the various rates of HSC division, differentiation and death were considered constant. The data collective comprise our previously published data from about Busch et al. (2015) and new data from another cohort of similar size, yielding a total of 180 fate-mapped mice, complemented by absolute cell counts for these populations (in one hindleg), and the BrdU pulse labeling data by Oguro et al. (2013) (Figure 3A; the subdivision of HSCs into HSC1 and HSC2 and of MPPs into MPP1-3 will be considered further below). Using maximum likelihood estimation, we found that the mathematical model (Methods) simultaneously fit the data on propagation of fate-mapping label (Figure 3B), on HSC numbers, which more than doubled from 6 to 36 weeks of age (Figure 3C), and on BrdU incorporation (Figure 3D). The best fit was obtained when symmetric differentiating HSC divisions were absent (*ρ* = 0), which indeed have not been observed *in vitro* (Loeffler et al., 2019). We found that 2.8-4.6% of HSCs per day divided symmetrically (95% confidence bounds of best fit value), which is the same magnitude as previous estimates of overall HSC proliferation (Oguro et al., 2013), whereas asymmetric divisions were inferred to occur at a much lower rate, 0-0.6% per day (Figure 3E).

**Figure 3.**
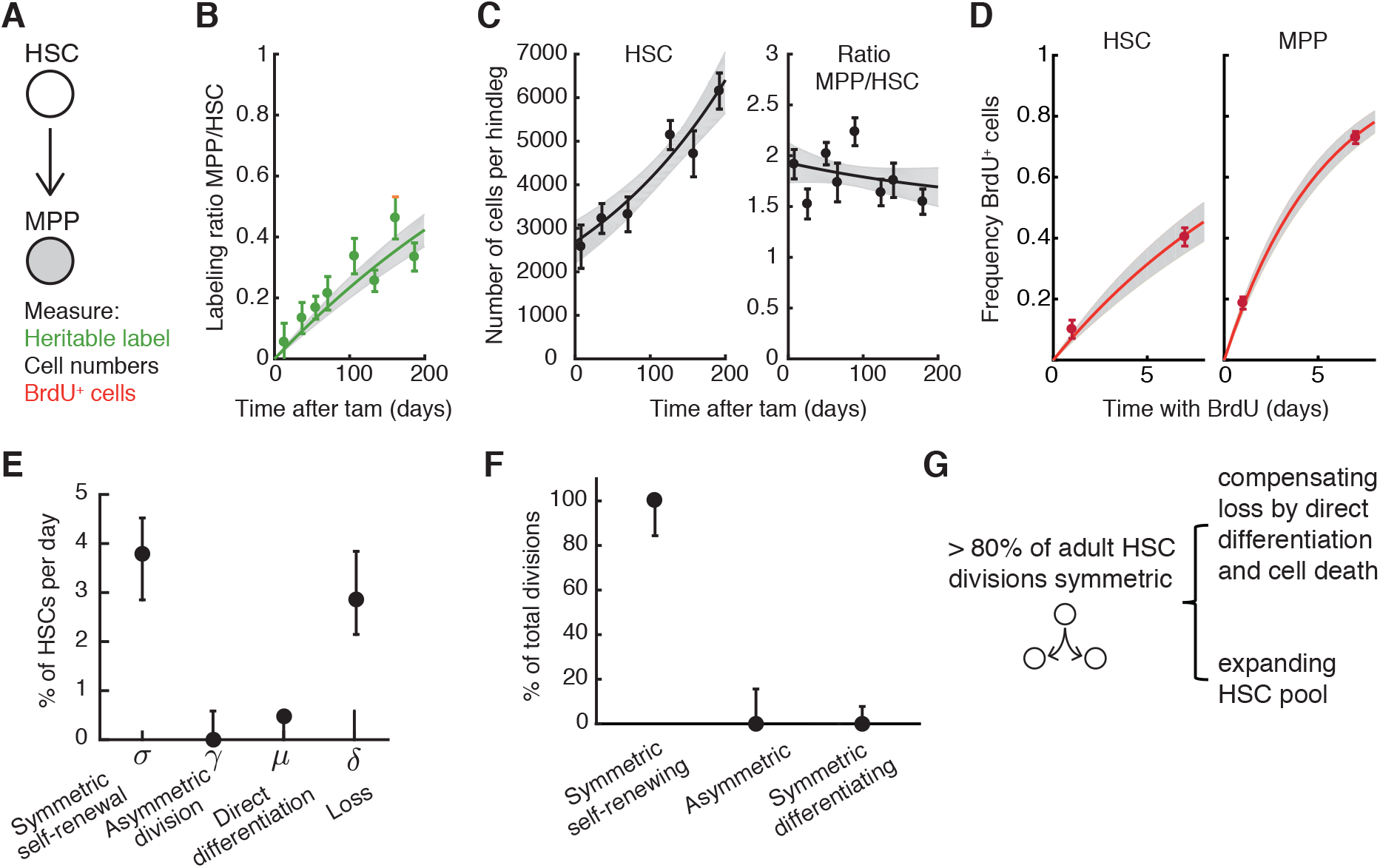
HSCs self-renew symmetrically. (A) Schematic of data used for parameter inference. (B) Model fit to the time course of fate-mapping label propagation from HSCs to MPPs, induced in *Tie2*^*MCM*^*Rosa26*^*YFP*^ mice 6-8 weeks of age (*N* = 12-49 mice per time points; error bars, SEM). (C) Model fit to HSC number in one hindleg and number ratio of MPP to HSC (*N* = 11-45 mice per time point; error bars, SEM). (D) Model fit to BrdU incorporation data taken from Oguro et al. (2013) (*N* = 5 mice per time point; error bars, SEM). (E) Inferred rates of HSC fates (dots, best fit; bars, 95% confidence bounds. (G) Inferred fractions for types of HSC divisions (dots, best fit; bars, 95% confidence bounds). (F) Symmetric HSC divisions supply cells for differentiation into MPPs, compensation of HSC death and expansion of the HSC population with age.

Up to 0.5% of HSCs per day underwent direct differentiation. The estimated HSC death was 2-4% per day, consistent with experimental data suggesting that HSC die by apoptosis *in vivo* (Domen et al., 2000). However, this rate may also include loss of HSCs when they spawn differentiating progeny that does not pass through an MPP stage (which is not counted as differentiation in the present model), as has been proposed for megakaryopoiesis (Rodriguez-Fraticelli et al., 2018; Carrelha et al., 2018).

As symmetric self-renewal appears to be the predominant division type of HSCs, we computed direct statistical estimates for the fractions of the different types of HSC divisions. At least four out of five HSC divisions are inferred to be symmetric self-renewal (Figure 3F). To understand how much of this self-renewal causes the increase in HSC number (Figure 3C), we estimated the exponential growth rate of the latter to be 0.42 % per day (doubling time of 165 days). Hence 9-15 % of the symmetric HSC divisions contributed to the expansion of the HSC population. The large remainder compensates HSC loss by direct differentiation and cell death (Figure 3G).

### Population output of HSCs to MPPs persists during aging

Next, we quantified changes of HSC differentiation with age. To this end, we included fate mapping data from older *Tie2*^*MCM*^*Rosa26*^*YFP*^ mice (label in HSCs induced between 6 and 8 weeks of age), up to 2 years of age (adding data from another 122 mice), and allowed the total differentiation rate per HSC, *α*_HSC_, to be a function of time. To determine *α*_HSC_, we estimated from the experimental data, at different time points, the temporal gradient in label acquisition by MPPs and the difference between label frequencies in HSCs and MPPs (Figure 4A). Factoring in the measured cell number ratio of MPPs to HSCs, these data allowed us to infer the function *α*_HSC_(*t*): The per-cell output of HSCs to MPPs significantly declined in older mice (Figure 4B). The differentiation rate appeared to peak intermittently in the first year (i.e., around 250 days) of life before declining, but the large confidence bounds for this time point preclude a definite statement. To characterize the overall decline of HSC differentiation rate with age, we fitted an exponential decay, which yielded a half-life of 9 months.

**Figure 4.**
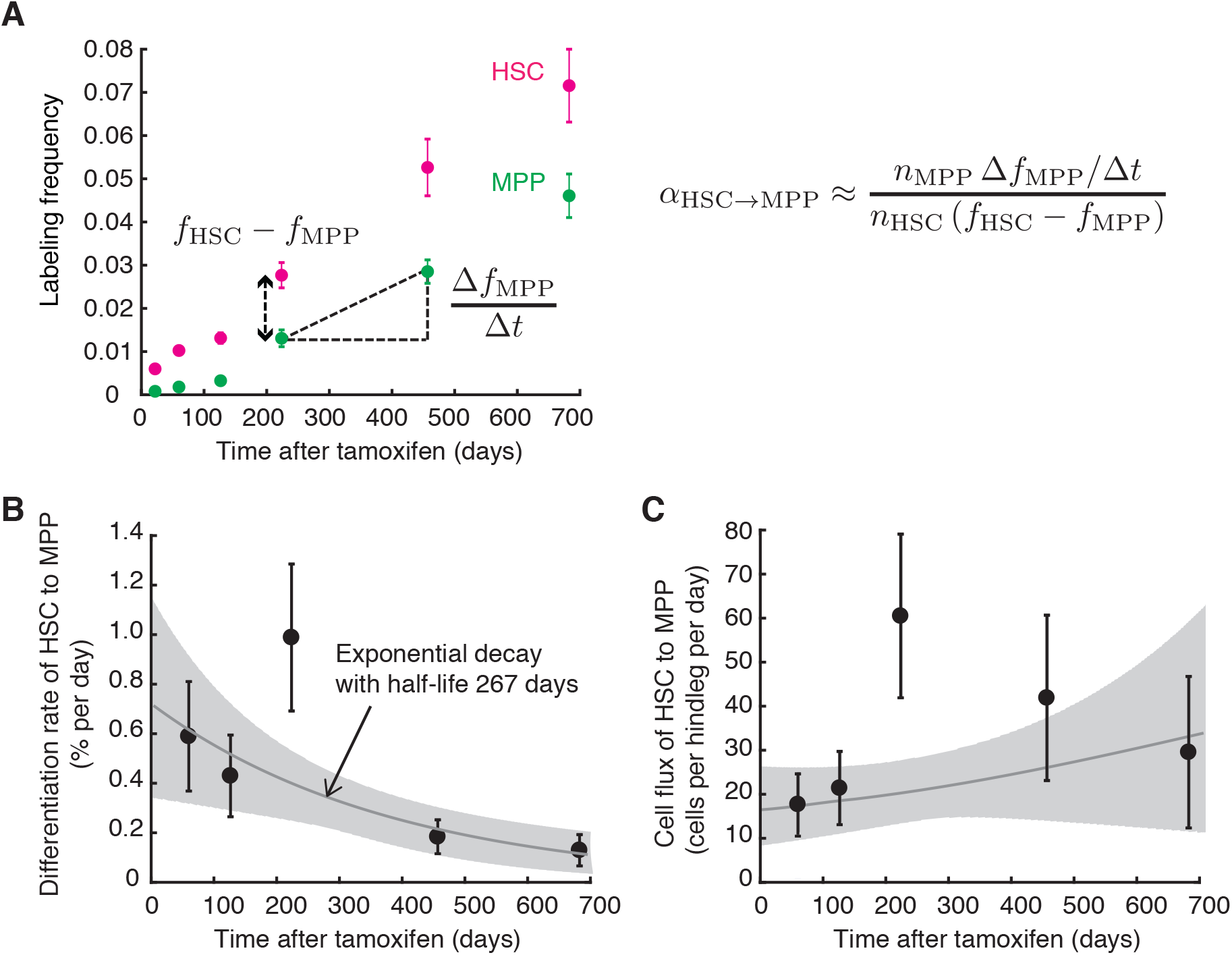
HSCs supply MPPs throughout the lifetime of the mouse. (A) Direct estimation of the time-dependent HSC differentiation rate from the fate mapping data. HSC label was induced in *Tie2*^*MCM*^*Rosa26*^*YFP*^ mice 6-8 weeks of age, and label frequency was measured in HSC and MPP populations up to 100 weeks later. *N* = 23-71 mice were pooled per time point for accurate determination of the mean error bars, SEM). Symbols in equations: *α*_HSC⟶MPP_, differentiation rate from HSC to MPP; *t*, time; *f*_*i*_, labeling frequency of population *i*; *n_i_*, cell count of population *i* (*i* = HSC, MPP). (B) Time-dependent output rate per HSC estimated from fate-mapping label propagation (dots, direct estimates from the experimental data; bars, 95% confidence bounds determined via non-parametric bootstrapping). The fit of exponential decay to the differentiation rate is also shown (grey line, best fit; shaded area, 95% confidence band). (C) Estimated cell flux from HSCs to MPPs obtained by multiplying the per-cell output rate with HSC number. The fit (grey line, best fit; shaded area, 95% confidence band) is based on the exponential decay fitted to the data in B and an exponential fit to HSC number over time (not shown).

While output per HSC declined, total HSC number continuously increased (Figure 3C), consistent with previous reports (Morrison et al., 1996; Sudo et al., 2000; Säwen et al., 2018). As a result, the total flux of cells from HSCs to MPPs, obtained as differentiation rate times HSC number, was rather age-independent as it never fell below the values reached in young mice (Figure 4C). The bone marrow extracted from the hindlegs contained between 2000 and 3000 phenotypic HSCs in young mice, thus comprising approximately 10-20 % of the total HSC number (Busch et al., 2015). Assuming that the inferred HSC differentiation rate can be extrapolated to other bone marrow locations, a total of more than 100 HSCs will turn into MPPs per day, regardless of age. These data indicate that, while the per-cell differentiation rate of HSCs declines slowly with age, the increase in HSC number fully compensates for this decline to maintain a small but steady supply of MPPs from HSCs throughout life.

### Transcriptome landscape shows lineage branching downstream of MPPs

To investigate when lineage differentiation sets in downstream of HSCs, we analyzed the transcriptional landscape of hematopoietic stem and progenitor cells (HSPCs). Much previous work on murine or human hematopoiesis has focused on progenitor cell transcriptomes, index-sorting specific progenitors (Paul et al., 2015; Olsson et al., 2016) or a broad population of HPSCs (Nestorowa et al., 2016; Velten et al., 2017; Giladi et al., 2018; Karamitros et al., 2018). A minority of cells in such HSPC samples will be HSCs which may under-represent them in the transcriptional landscape. To analyse a balanced representation of stem and progenitor cells, we sorted separately HSCs, MPPs, HPC1 and common myeloid progenitors (Lin^−^Kit^+^Sca1^−^CD16/32^low^ CD34^+^), subjected these populations to single-cell RNA sequencing using the 10X Chromium platform, and thus obtained ~2000-3000 single-cell transcriptomes per population after quality control. We reasoned that differentiating cells will undergo gradual transcriptional changes when progressing towards progenitor stages. To identify the resulting differentiation trajectories in transcript space, we used the diffusion map, which preserves both local neighbors and overall nonlinear differentiation trajectories in lower-dimensional representations of the data (Haghverdi et al., 2016). This analysis revealed a branched transcriptional landscape (Figure 5A; shown are two-dimensional projections in the first three diffusion components). The stem consists of HSCs followed by, and partially overlapping with, MPPs. Downstream of the stem, branches occur in HPC1 and CMPs. The fraction of cycling cells was similar in HSCs and MPPs but increased in HPC1 and further in CMPs, as measured by signature genes for S, G2 and M phases of the cell cycle (Figure 5B). Hence single-cell RNA-seq indicates that differentiation events in the transcriptome occur downstream of the common stem.

**Figure 5.**
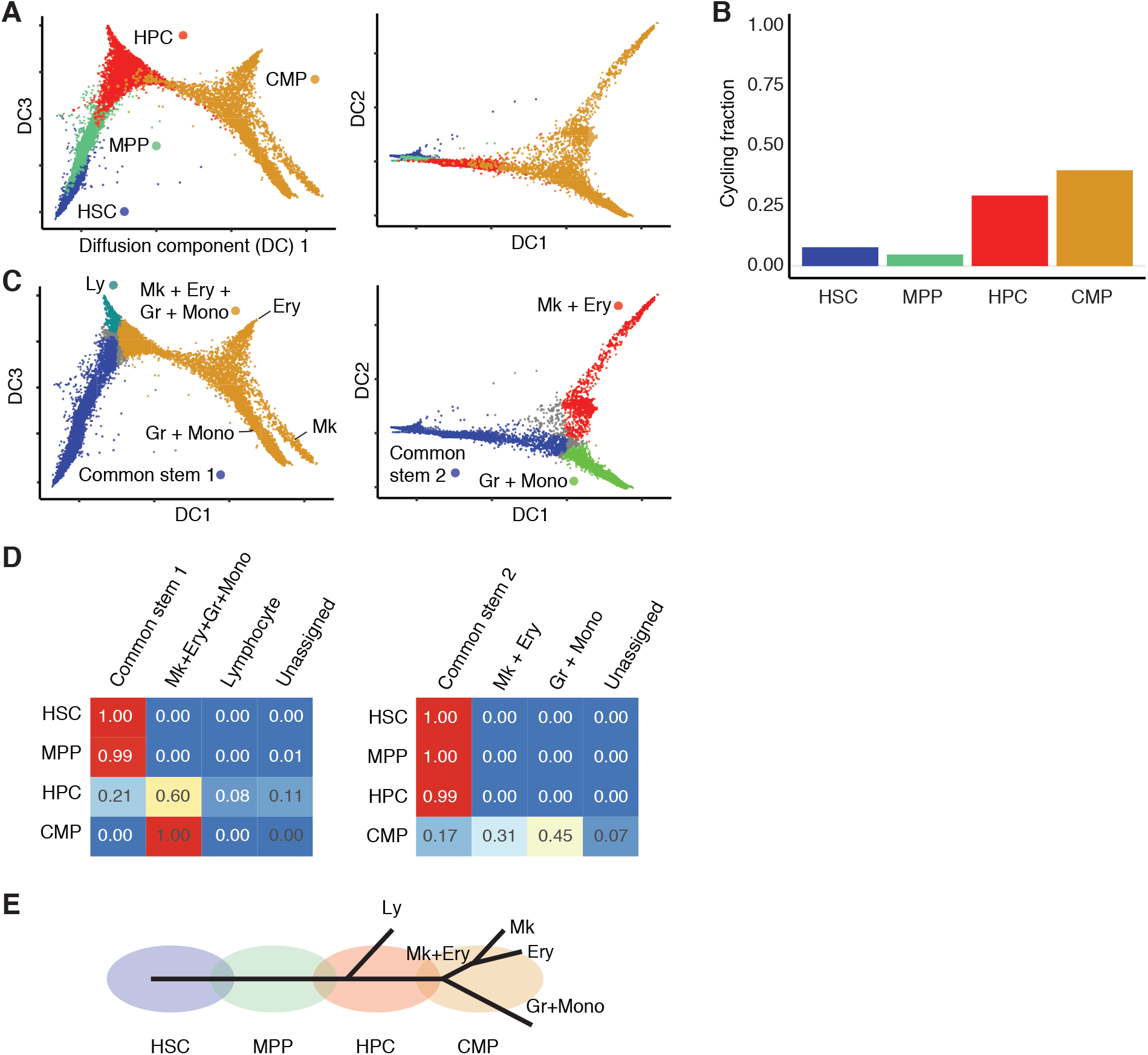
Joint analysis of single-cell transcriptomes oh HSCs, MPPs, HPC1 and CMPs. (A) Diffusion map representation of single-cell transcriptomes obtained from sorted HSC, MPP, HPC1 and CMP populations. Projections in the first three diffusion components are shown. (B) Fractions of cycling cells (with S or G_2_/M gene signatures) in the respective populations. (C) Branch analysis using diffusion pseudotime detects major branching events within the HPC1, splitting of lymphocyte (Ly) differentiation and CMP populations, with a bifurcation between cells with megakaryocyte (Mk)-erythroid (Ery) and granulocyte (Gr)-monocyte (Mono) signature; cells that could not unambigously assigned are labeled in grey. (D) Fraction of cells assigned to distinct branches by diffusion pseudotime analysis; left, with respect to lymphocyte branching in HPC1; righ, with respect to the split between Mk-Ery and Gr-Mono branches in CMP. (E) Schematic of the branching structure of the transcriptional landscape.

We annotated the transcriptomes of individual cells using known lineage markers (Figure 5C) and employed the diffusion pseudo-time algorithm (Haghverdi et al., 2016) to map cells to individual branches (Figure 5D). All HSCs and MPPs were assigned to the common stem. Within HPC1, a small fraction of cells were mapped together with all HSCs and MPPs to the common stem, while the large remainder were placed beyond the first branching event, with 8% of assignable HPC1 showing a lymphoid signature and 76% a CMP-type signature (Figure 5D, left panel). This finding is consistent with single-cell functional analyses of HPC1, showing that this population contains a heterogeneous collection of restricted progenitors (Oguro et al., 2013). The next branching of the transcriptome landscape occurred within the CMPs, separating cells with a granulocyte-monocyte transcript signature from cells with megakaryocyte-erythroid signature (Figure 5D, right panel). The latter sub-branch split further into megakaryocyte and erythroid branches.

The transcriptional landscape of HSCPs suggests the following features of unperturbed hematopoiesis: (1) Tip stem cells give rise to gradually differentiating cells, as indicated by the common stem of HSCs and MPPs (Figure 5E); (2) Differentiation into lymphoid and myeloid lineage branches becomes apparent by differential gene expression in populations downstream of MPPs; and (3) At approximately the same stage at which lineage differentiation becomes manifest in the transcriptome, cell proliferation increases (cf. Figure 5B).

### HSCs and early-stage MPPs self-renew in physiological hematopoiesis

We examined the key predictions derived from the analysis of the HSPC transcriptome landscape by fate mapping. The first prediction, gradual differentiation from HSC, is consistent with the results of fate mapping, indicating the progression HSC1 ⟶ HSC2 ⟶ MPP1+2 ⟶ MPP3 ⟶ HPC1 (see Figure 1C). To gain quantitative insight, we modeled propagation of fate-mapping label through this differentiation pathway (Figure 6A). To this end, we assumed that HSC1 preferentially contained tip HSCs that are responsible for the increase in labeling frequency in aging mice (cf. Figure 1D). This model provided an excellent fit to a large collection of experimental data on label propagation (Figure S1A and S1B), cell numbers (Figure S1C-S1E) and BrdU incorporation (Figure S1F). Of note, the model predicted the labeling frequency in tip HSC1 to be at least 12% (Figure S1G), which could be tested by crossing *Tie2* reporter mice (Ito et al., 2016) with *Tie2*^*MCM*^ fate mapping mice (Busch et al., 2015). Moreover, the model correctly predicted cell number data that were not used for fitting (Figure S1H), thus supporting its validity.

**Figure 6.**
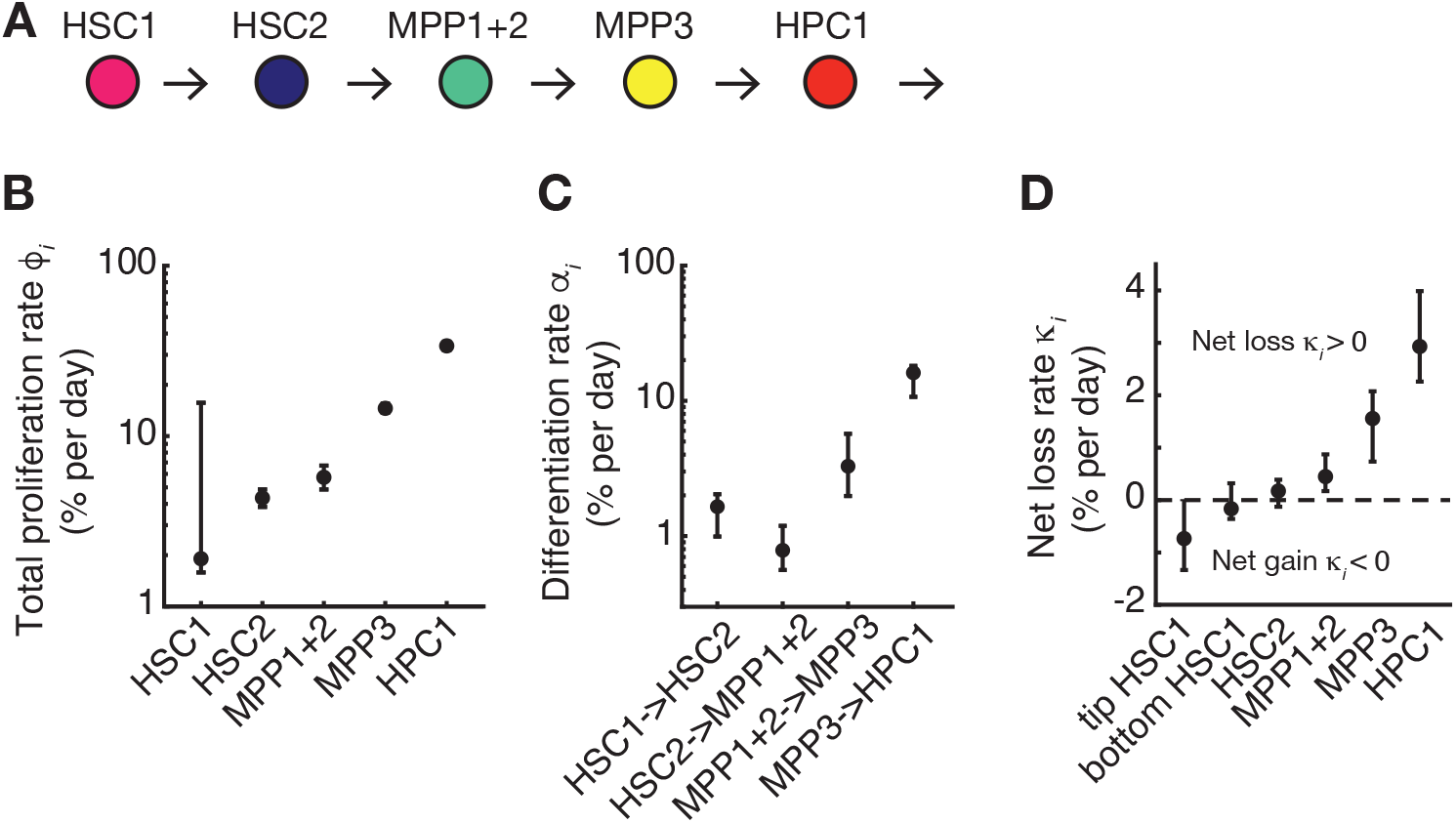
Quantification of fates of hematopoietic stem and progenitor cells in young adult mice. (A) Linear differentiation pathway indicated by fate mapping (see Figure 1C). (B) Total cell proliferation rates. (C) Differentiation rates of precursors to their respective products. (D) Net loss rates. The data in (B)-(D) result from fitting the linear differentiation model (A) simultaneously to the fate mapping and BrdU incorporation data (cf. Figure 1C,E). Dots, best fit values; error bars, 95% confidence intervals. Note that the majority of inferred rates have narrow confidence intervals, implying that the combined fate mapping and BrdU incporporation data are informative on these parameters.

Inferring the total proliferation rate, we found a very small best fit for HSC1 (~ 2% per day), while the statistical uncertainty of this estimate was large (Figure 6B). More certain inferences were obtained for HSC2 and MPP1+2, which had similar proliferation rates in the range of 3-5% per day. In downstream populations (MPP3, HPC1), proliferation rate was strongly increased, agreeing also with previous data on loss of H2B-GFP fluorescence (Oguro et al., 2013). In sum, the inference of HSPC proliferation rates from labeling data (Figure 6B) is consistent with the observed increase in the fraction of cycling cells when progressing from MPPs to HPC1 in the transcriptome landscape (cf. Figure 5B).

Concomitant with the rise in proliferation rate, the differentiation rate also increased strongly for the transition from MPP3 to HPC1 (Figure 6C); about 1% of HSC2 differentiated to MPP1+2 per day whereas ~15% of MPP3 gave rise to HPC1. Importantly, the two estimates for the differentiation from HSCs to MPPs obtained by lumping HSC1 and HSC2 as well as all MPP subpopulations (Figure 4) and by quantifying the more resolved HSC2⟶MPP1+2 transition at the boundary of the two populations overlapped, indicating that a finer level of subpopulation resolution does not strongly alter the inferred differentiation rates. In sum, the onset of lineage differentiation seen in the transcriptional landscape coincides with increases in both progenitor cell proliferation and differentiation rates.

To understand to what extent the different stem and progenitor populations compensate cell differentiation and loss by self-renewal or rely on input from upstream populations, we inferred the net loss rates *κ* (Figure 2B). The estimated numbers (Figure 6D) indicate that the continuous expansion of the HSC compartment as a whole is fuelled by the tip-HSCs within HSC1; indeed these cells have a net gain (i.e., negative net loss rate). This finding indicates that tip HSCs exhibit excess self-renewal. Up to MPP1+2, the net loss rates stay close to zero, implying that self-renewal in these cell populations is sufficient to maintain the output of more differentiated cells over very long time periods, with minimal input from their respective precursors. By contrast, MPP3 and HPC1 had sizeable positive net loss rates, implying that these populations are more dependent on input for their maintenance than all upstream compartments. Taken together, these data suggest that self-renewal in physiological hematopoiesis extends beyond the HSCs (HSC1+HSC2, which have previously been assigned as stem cells by transplantation experiments), and includes MPP1+2 (which in transplantation are only short-term repopulating).

### MPPs slowly contribute to granulopoiesis

In the single-cell transcriptome data, lineage branching becomes evident downstream of MPPs. To shed light on the differentiation fluxes towards the major lineages, granulocytes, erythrocytes, megakaryocytes and lymphocytes we analysed propagation of fate-mapping label towards these lineages (Figure 7A; these data were measured in the same mice that so far have been analysed for fate mapping in HSPCs). According to the fate-mapping inequality (Box 1), the labeling frequency in a cellular product is smaller than, and becomes at most equal to, the labeling frequency of its precursor. This inequality was satisfied for all mature cell populations relative to HSCs and MPPs (Figure 7B, blue rectangles; note the shorter time period for measuring erythrocyte progenitors). For all cell types, except lymphocytes, it was violated at least at one time point in HPC1, and for megakaryocytes also in CMPs (Figure 7B, red asterisks). The single-cell transcriptome data suggest an explanation for this apparent violation of the fate-mapping inequality, as follows: Within HPC1, lymphoid differentiation branches off. If lymphoid differentiation is slower than myeloid (which is likely the case as judged by slow label accumulation in B cells versus granulocytes), lymphoid biased HPC1 will carry less fate-mapping label than HPC1 with myeloid fate and hence the overall labeling frequency in HPC1 stays below that of the myeloid differentiation pathway. The same argument may apply to erythropoiesis and megakaryopoiesis relative to lymphopoiesis. Hence lineage branching within phenotypic progenitor populations in the transcriptional landscape (Figure 5B, D) is reflected in an apparent violation of the fate-mapping inequality, which indicates that the different fate-biased subpopulations in HPC1 and CMPs also have different kinetics. However, no such kinetic heterogeneity is evident in the MPPs. Indeed, HSC fate mapping data is consistent with MPPs being precursors of granulocytes, erythrocytes, megakaryocytes and B lymphocytes.

**Figure 7.**
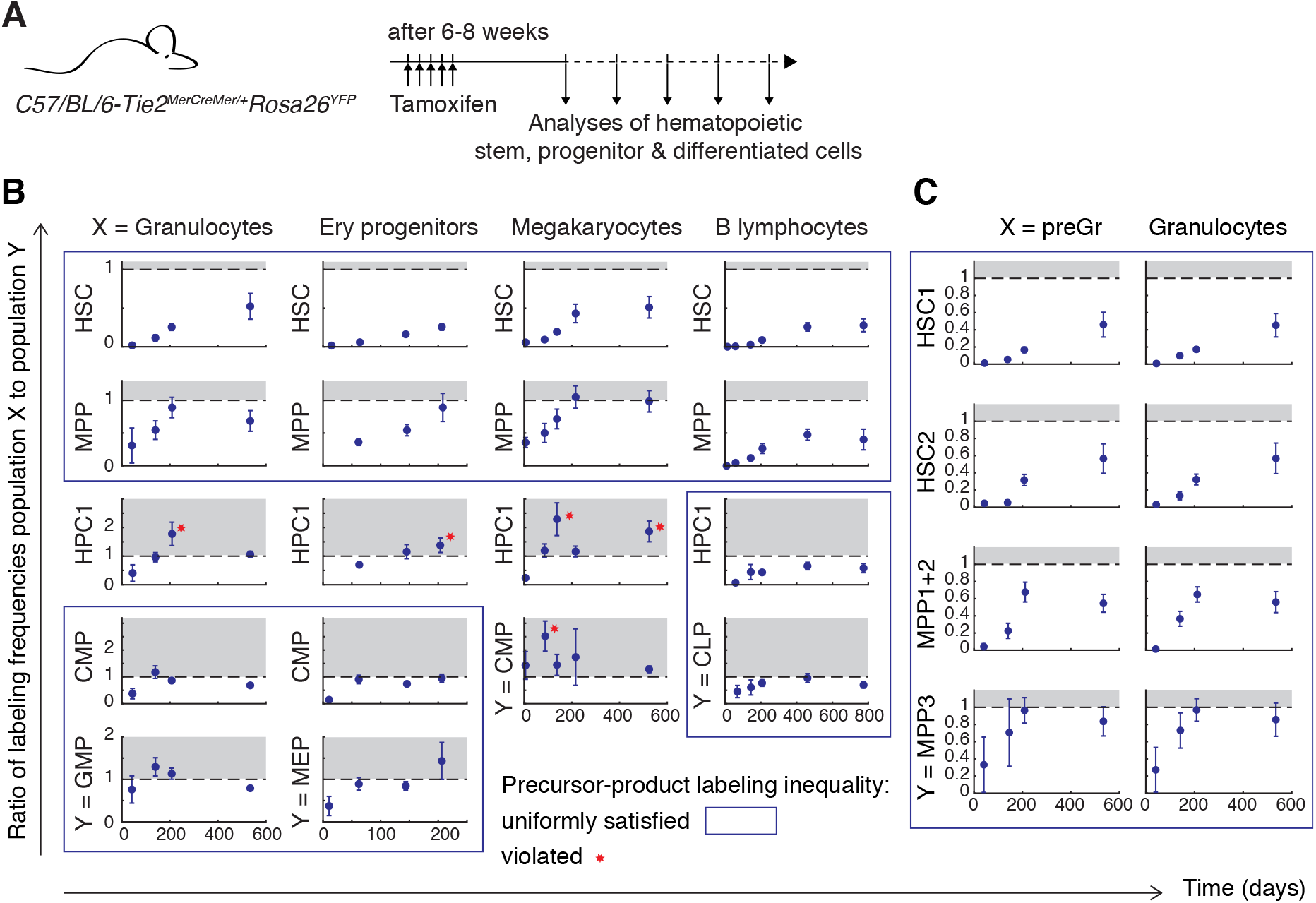
Label equilibration of mature cell populations with MPPs. (A) Scheme of the experiment; for details see Figure 1. (B) Labeling frequencies in mature population relative to labeling frequencies in the indicated stem and progenitor cell types. (C) Finer resolution of granulocyte labeling frequencies relative to labeling frequencies in HSC and MPP subpopulations. Blue frames indicate that fate mapping inequality for precursor-product pairs is satified; red stars indicate data that show significant violation of this inequality. The respective precursor population cannot be a uniform source of the respective mature product.

Granulocytes, erythrocytes and megakaryocytes make up the major part of hematopoietic cell production in adult mice (Mackey, 2001). In agreement with this, granulocytes, erythrocyte progenitors and megakaryocytes equilibrated their label with MPPs much faster than B lymphocytes, which did not reach equilibration over the lifetime of the mouse (Figure 7B). Nevertheless, label equilibration for these mainstay outputs of hematopoiesis occurred only about 200 days after label induction in HSCs (note the different time axis for erythrocyte progenitors in Figure 7B). To study this phenomenon at higher resolution, we focused on granulocytes and evaluated labeling ratios with HSC1, HSC2 and MPP1-3. We found rapid label equilibration with MPP3, but no equilibration with upstream populations (Figure 7C). This finding echoes the abrupt acceleration of cell differentiation in MPP3 compared to upstream HSPCs (cf. Figure 6C) and indicates that MPP3 quantitatively supply granulopoiesis. Nevertheless, label equilibration of granulocytes with MPP3 still took about 200 days. These data indicate that progenitor populations downstream of MPP3 are replenished only slowly over about half a year and hence are predicted to behave as medium-term sources of cell production, rather than just transit amplifiers under steady-state conditions.

## DISCUSSION

Here we developed an inference framework for the *in vivo* dynamics of stem cells that integrates data from stem cell fate mapping by heritable labeling with data reporting on cell-cycle-dependent label incorporation. Combining these two independent cell-population labeling techniques provides insight into decisions of stem cells that have so far been inaccessible by using these approaches separately (van der Wath et al., 2009; De Boer and Perelson, 2013; Busch et al., 2015). The fact that the mathematical model hold for any type of time-dependent dynamics and can accommodate time-varying rate parameters makes this quantitative framework applicable to studying changes in stem cell decisions over the course of life, as done here, and also stem cell responses to injury or disease (Rulands and Simons, 2016).

Our results contribute to the understanding of physiological hematopoiesis in several respects. We infer that HSCs divisions are predominantly symmetric self-renewing during unperturbed physiological hematopoiesis. While HSCs in cell culture are stimulated to divide more frequently than in unperturbed hematopoiesis *in vivo*, Loeffler et al. (2019) observed 80-85% of divisions to be symmetric *in vitro*; assigning symmetry or asymmetry based on lysosome inheritance, which was related to subsequent fate (Loeffler et al., 2019). In contrast to these data, another *in vitro* study, judging asymmetry by distribution of the Rho GTPase Cdc42, reported preferential asymmetric divisions of HSCs from young mice, and symmetric divisions of HSCs from older mice (Florian et al., 2018) (interestingly, this behavior was the opposite of increasing asymmetry with age observed with neural stem cells (Bast et al., 2018)). Our *in vivo* estimate of 80-100% symmetric divisions agrees with the *in vitro* data by Loeffler et al. (2019), suggesting that HSCs may have an intrinsic propensity for symmetric self-renewal. Consistent with predominant symmetric self-renewal, we observed an increase of HSC number already in young mice, without enhanced differentiation output per cell, which extends previous data reporting such an increase in aged mice (Morrison et al., 1996; Sudo et al., 2000).

Our rate estimates also show that a large part of HSC self-renewal compensates for HSC loss. This loss can occur by direct differentiation or cell death. Compared to the 3-5 % symmetric self-renewal per day, we found that about 1% of HSCs per day differentiate into MPPs in young mice, quite independent of whether we employed the usual (HSC⟶MPP) or a more resolved (HSC2⟶MPP1+2) definition of this transition, attesting to the robustness of experimental data and inference. Moreover, any MPP-independent differentiation, for example into megakaryocytes (Rodriguez-Fraticelli et al., 2018; Carrelha et al., 2018), as well as loss by death, will together be in the order of 2-3 % per day. HSC death by apoptosis is likely to occur at appreciable rate, as implied by the 2-fold increase in HSC numbers upon overexpression of the anti-apoptotic protein Bcl-2 (Domen et al., 2000). Taken together, our findings are in line with the early conjecture by Lajtha et al. (1962) that HSC numbers are set by the balance of symmetric self-renewal with differentiation and death.

A remarkable finding of our approach is that the low differentiation rate of HSCs into MPPs, of a few hundred cells per day in the mouse, is maintained over the entire lifetime, despite a decline in HSC differentiation rate. As hypothesized previously (Säwen et al., 2018), the compensatory mechanism is a steady increase in HSC number, which we show here to be driven by tip HSCs enriched within HSC1. Following Busch et al. (2015), there have been several fate mapping studies of hematopoiesis in mice that appeared to argue that HSC output rate to MPPs is much larger (Sawai et al., 2016; Chapple et al., 2018; Säwen et al., 2018). However, these experiments are not directly comparable to ours, as the Cre drivers and induction conditions used caused initial labeling of HSCs together with progenitor cells, naturally causing the label to appear more rapidly in downstream cell populations than after selective HSC labeling via *Tie2*^*MCM*^ reported here and previously in Busch et al. (2015). As we have shown here, mathematical inference can disentangle the contributions of multiple labeled sources (in this paper: HSC1 and HSC2), provided that the upstream source has initially higher labeling frequency. With a similar approach, Sawai et al. (2016) found an HSC-to-MPP differentiation rate that is of the same order of magnitude (2.6 % of HSCs per day differentiating into MPPs) as our estimate, and hence does not contradict the idea of infrequent HSC input to MPPs. The remaining quantitative difference to a rate of ~ 1 % per day, and lower in old mice, estimated from our data, could be due to the fact that *Tie2*^*MCM*^ labels more selectively tip HSCs than the Cre driver used by Sawai et al. (2016), and might also be affected by the serial bone marrow aspiration employed in that study, which could stimulate HSC differentiation. Moreover, Sawai et al. (2016) have questioned whether *Tie2*^*MCM*^-driven fate mapping quantitatively supplies hematopoietic lineages. Here we have given an affirmative answer by showing that the HSC population reached by this Cre is the unique upstream source of MPPs. This conclusion is independent of the initial HSC labeling frequency.

To understand how cells proceed to differentiation when leaving the HSC state, we juxtaposed label propagation with single-cell transcriptomes. The key features of these independent sets of data are remarkably concordant. In particular, label propagation implies that populations from tip HSCs to early-stage MPPs (MPP1+2) self-renew. In the transcriptome landscape, the corresponding cells are located in the common stem, where cells do not express lineage gene modules. Kinetically, differentiation and division rates of HSC1 through MPP1+2 are small, up to a few percent per day, consistent with a small fraction of cycling cells in the common stem. Taken together, these data suggest that the common stem of the transcriptional landscape, including early-stage MPPs, consists of stem cells, of which only a small fraction is in cell cycle or proceeds to differentiate at any given time. Within this extended stem cell population, there is still a hierarchy, as tip HSC have excess self-renewal and drive slow expansion of the stem cell population as a whole. The transition of cells to differentiation is accompanied by increases of proliferation and differentiation rates and a drop of self-renewal, which become significant in late-stage MPPs (MPP3), just before the first lineage bifurcations occur in the transcriptional landscape. The transcriptional identification of those rare cells undergoing differentiation at the time of cell isolation for single-cell RNA-seq remains elusive.

Recent work, linking HSC barcoding and single-cell transcriptomics, has provided transcriptional signatures for HSC clones with multilineage (myeloid, erythroid and lymphoid) output and myelo-eyrhroid restricted output (Pei et al., 2020), as well as for differentiation-inactive HSC clones (Bowling et al., 2020; Pei et al., 2020). Importantly, differentiation-inactive HSC clones are not proliferatively quiescent (Pei et al., 2020), which also supports our finding of symmetric self-renewal of HSCs.

We have shown here that downstream populations, including early-stage MPPs, are capable of self-renewal, and HSC ablation experiments have indicated that hematopoiesis remains normal for more than six months practically without HSCs (Schoedel et al., 2016; Sheikh et al., 2016); also note that we find here that granulopoiesis is supplied by progenitors downstream of late-stage MPP for several months. So why is the small output rate of tip HSCs maintained for the lifetime of the mouse? We suggest that the trickling down of cells generated by tip HSCs limits the lifetime of existing MPP clones. These less-divided cells will spawn new MPP clones that replace older ones in which MPPs have already divided more and, accordingly, accumulated somatic mutations. Therefore, tip HSCs in adult mice could function as a source of little damaged cells that found new hematopoietic clones throughout the life of the organism. This mechanism would be akin to low-level input of thymic progenitors prevents leukemia by regularly outcompeting thymus-resident progenitors (Martins et al., 2014; Gao et al., 2019). The recent findings that hematopoiesis in the much more long-lived humans tends to become clonal with age (i.e., derive to a considerable extent from few stem cells that spawn very large clones) and that clonal hematopoiesis increases leukemia risk (Bowman et al., 2018) support the notion that mutation accumulation in HSCPs is a serious risk, which is normally controlled by slow differentiation of rarely dividing tip HSCs.

## Acknowledgments

We thank Sean Morrison and Hideyuki Oguro (University of Texas Southwestern Medical Center, Dallas) for providing the raw data of BrdU incorporation; Nina Claudino (DKFZ) for expert technical assistance and all members of the Höfer and Rodewald groups for ongoing support and discussions. Support from DFG (SFB873-B11 to HRR and TH; Leibniz program to HRR), the Helmholtz Association II program (to HRR), ERC (Advanced Grant 742883 to HRR) and EU-FP7 (317040, QuanTI, to TH) is gratefully acknowledged.

## Box 1: Precursor-product rules in fate mapping

The mathematical law for the propagation of fate-mapping label (Materials and Methods) has general implications for the interpretation of fate mapping data. We consider a fate mapping experiment where a permanent source of fate-mapping label is switched on in a stem cell population at *t* = 0 (realized here by tamoxifen administration triggering nuclear import of Cre expressed in HSCs). Ideally, no other population would initially be labeled (e.g., if Cre expression were highly specific to stem cells). However, for the following rules of fate mapping to hold, there could be some initial labeling of downstream populations, provided that the initial labeling frequency decreases monotonically with developmental distance from the stem cells. For example, in the experimental data shown in Figure 1C, HSC1 are the tip stem cell compartment and most strongly labeled initially, while HSC2 are labeled more weakly, and progenitors are not labeled.

We analyse the labeling frequencies in a pair precursor and product populations, P and Q, respectively:

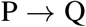

P could be the tip stem cells or any compartment downstream. Given the precursor labeling frequency over time, *f*_P_(*t*), the labeling frequency of the product, *f*_Q_(*t*), is related to it through the differential equation:

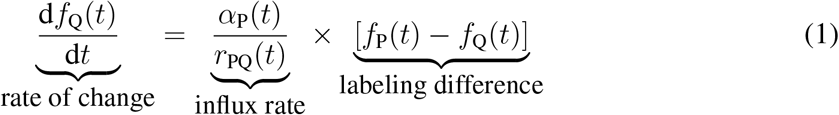

where *α*_P_(*t*) is the total rate of precursor differentiation to product and *r*_PQ_(*t*) is the ratio of product to precursor cell numbers. From this law of fate-mapping label propagation, two qualitative rules follow that can easily be checked with experimental data.

### Rule 1—Fate mapping inequality

Given that the labeling frequency of the product is smaller than that of its precursors at a time point *t*_0_, this remains so for all later times

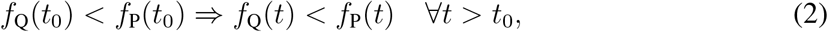

where the difference, however, can become infinitesimally small. This can be seen as follows. First, we take the precursor to be the tip stem cells with permanent labeling, such that *f*_P_(*t*) = *f*_P_(0) = const. Then, under the above condition of monotonically deceasing initial labeling frequencies away from the tip stem cells, we have *f*_P_(0) − *f*_Q_(0) > 0 and hence

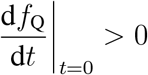

As a consequence, *f*_Q_ will increase and approach *f*_P_ from below. This argument can now be propagated downstream. Consider the previous products as the new precursors for one of its own product populations. Then at some *t*_0_ ≥ *t*, we again have *f*_P_(*t*_0_) − *f*_Q_(*t*_0_) > 0 and hence the labeling frequency of this product will also tend to equilibrate with its precursor from below. The key point is that at no time any of the labeling frequencies will decrease, which is guaranteed by the initial condition of preferential labeling of tip stem cells with monotonically decreasing labeling frequencies away from them.

Immunophenotypically defined populations (P or/and Q) could consist of subpopulations. For example, MPP can be subdivided into MPP1, MPP2 and MPP3; however, in general subpopulations may exist that cannot be distinguished by known cell surface markers. Even with such unidentified heterogeneity, the fate mapping inequality continues to hold if the subpopulations form a developmental sequence. To be specific, let us assume that both precursor and product consist of subpopulations:

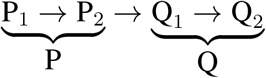

of which we only know the average labeling frequencies

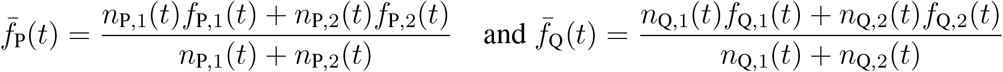

where, as before, the *n*_*i*_ denote total cell numbers in the respective subpopulations. From Inequality 3, we have

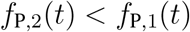

which implies

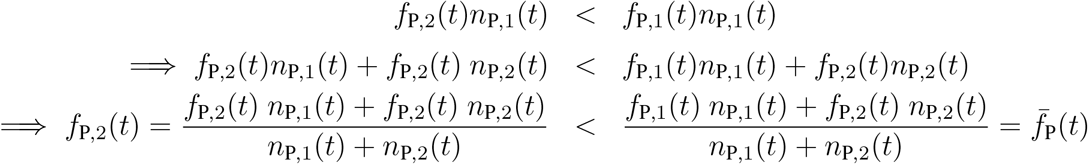

By analogous arguments, it can be established that

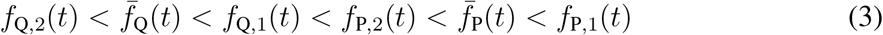

and hence the fate mapping inequality holds for the average labeling frequencies even if P and Q are heterogeneous.

### Rule 2—Equilibration with source

Equation 1 for label propagation from product P to precursor Q assumes that P is the unique source of Q. Over a sufficiently long period of time, the two labeling frequencies will equilibrate:

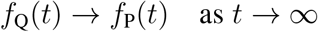

If both differentiation rate and ratio of product to precursor cell numbers are approximately constant, the characteristic time scale for equilibration is

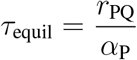

which is inversely proportional to the differentiation rate.

Fate-mapping label typically marks only a fraction of stem cells. This could be due to a probabilistic component in Cre action (Pei et al., 2017) or the unlabeled stem cell fraction could have inherently different properties from the labeled one (e.g., these cells might not express Cre). In the latter case, there could be two distinct precursor populations of Q: P_1_, deriving from the labeled stem cell population, and P_2_, deriving from the intrinsically unlabeled stem cell population.

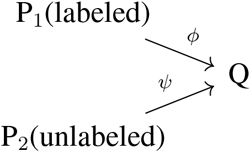

We denote the flux of cells from P_1_ to Q by *ϕ* = *α*_P,1_*n*_P,1_, and the flux from P_2_ to Q by *ψ* = *α*_P,2_*n*_P,2_. The unlabeled source effectively dilutes the label in Q, which is expressed in the following balance equation, generalizing Equation 1,

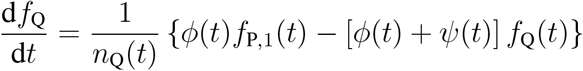

Consequently, the labeling frequency of the product approaches, for monotonic time dependencies of the fluxes,

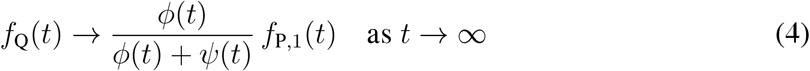

which remains smaller than the labeling frequency of the labeled precursor P_1_. Equation 4 is the general law of label equilibration in the product in the presence of labeled and unlabeled precursor populations. If in experimental data the labeling frequency of the product stays below that of the precursor, there could be two principal reasons: (i) there is an additional unlabeled precursor population, or (ii) label equilibration is too slow to be completed over the duration of the experiment. Therefore, no definite conclusion can be drawn from such data. However, if precursor and product labeling frequencies do equilibrate in an experiment then the labeled precursor population will be the unique source of the product. This conclusions holds irrespective of how large the labeling frequencies are in absolute terms.

---

**Figure S1.**
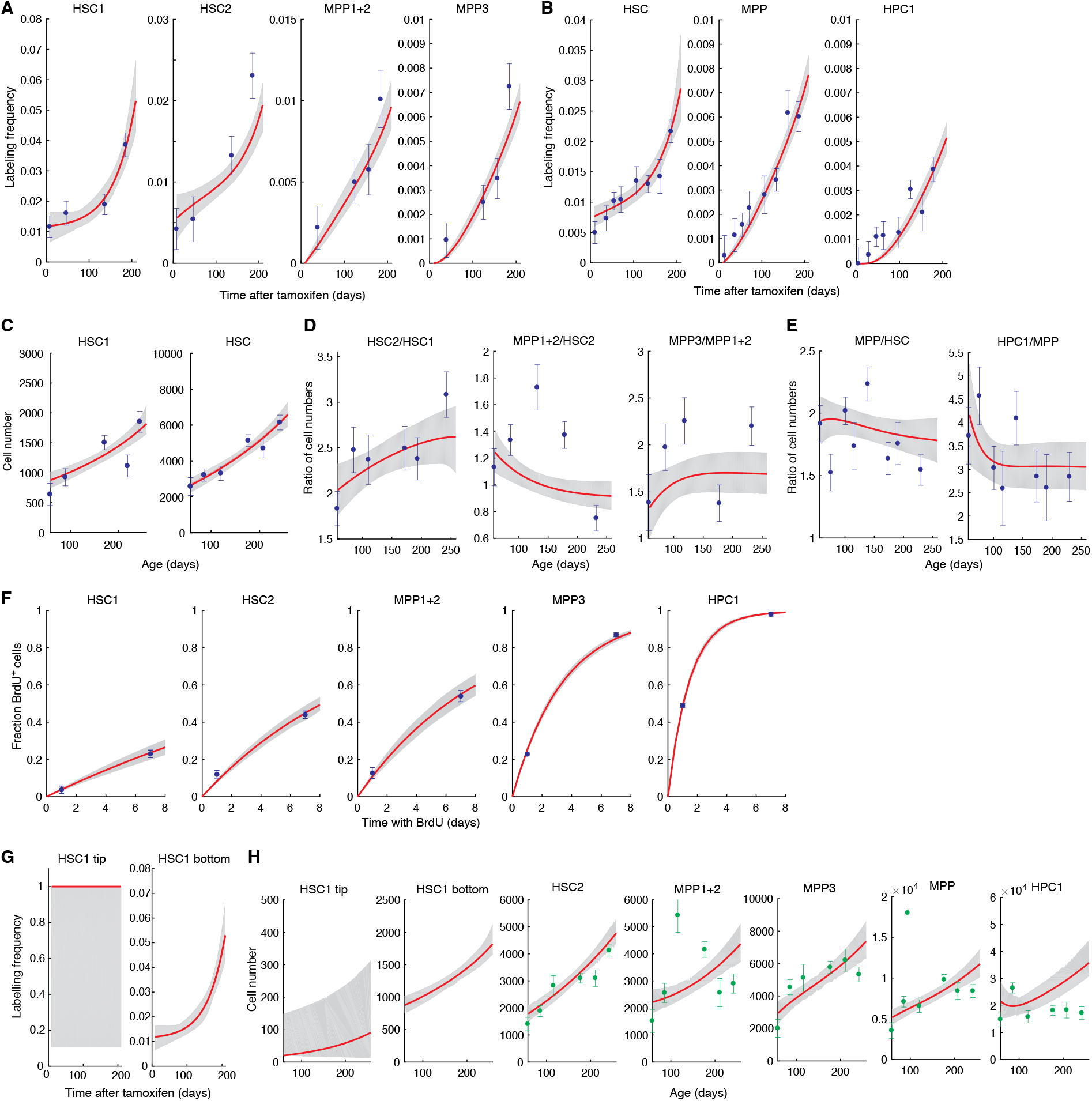
Fit of the model for HSPC dynamics to experimental data, related to Figure 6. (A) Model fit to the time course of fate-mapping label propagation in HSC1, HSC2 and MPP1-3, induced in *Tie2*^*MCM*^*Rosa26*^*YFP*^ mice 6-8 weeks of age (*N* = 11-18 mice per time points; error bars, SEM). (B) Model fit to the time course of fate-mapping label propagation in HSC, MPP and HPC1, induced in *Tie2*^*MCM*^*Rosa26*^*YFP*^ mice 6-8 weeks of age (*N* = 12-49 mice per time points; error bars, SEM). (C) Model fit to HSC1 and HSC numbers in one hindleg (for mouse numbers see D and E, respectively). (D) Model fit to cell number ratios for populations downstream of HSC1 (*N* = 11-18 mice per time points; error bars, SEM). (E) Model fit to cell number ratios for populations downstream of HSC (*N* = 12-49 mice per time points; error bars, SEM). (F) Model fit to BrdU incorporation data taken from Oguro et al. (2013) (*N* = 5 mice per time point; error bars, SEM). (G) Predicted labeling frequencies of tip HSCs in HSC1 and the remainder of HSC1 cells (“bottom” HSC1). (H) Predicted cell numbers. Where available, experimental data, which have not been used for model fitting, are shown.

## References

Akinduro, O., Weber, T. S., Ang, H., Haltalli, M. L. R., Ruivo, N., Duarte, D., Rashidi, N. M., Hawkins, E. D., Duffy, K. R. and Lo Celso, C. (2018). Proliferation dynamics of acute myeloid leukaemia and haematopoietic progenitors competing for bone marrow space. Nat Commun 9, 519.

Bast, L., Calzolari, F., Strasser, M. K., Hasenauer, J., Theis, F. J., Ninkovic, J. and Marr, C. (2018). Increasing Neural Stem Cell Division Asymmetry and Quiescence Are Predicted to Contribute to the Age-Related Decline in Neurogenesis. Cell Rep 25, 3231–3240.

Bernitz, J. M., Kim, H. S., MacArthur, B., Sieburg, H. and Moore, K. (2016). Hematopoietic Stem Cells Count and Remember Self-Renewal Divisions. Cell 167, 1296–1309.

Bowling, S., Sritharan, D., Osorio, F. G., Nguyen, M., Cheung, P., Rodriguez-Fraticelli, A., Patel, S., Yuan, W. C., Fujiwara, Y., Li, B. E., Orkin, S. H., Hormoz, S. and Camargo, F. D. (2020). An Engineered CRISPR-Cas9 Mouse Line for Simultaneous Readout of Lineage Histories and Gene Expression Profiles in Single Cells. Cell 181, 1410–1422.

Bowman, R. L., Busque, L. and Levine, R. L. (2018). Clonal Hematopoiesis and Evolution to Hematopoietic Malignancies. Cell Stem Cell 22, 157–170.

Busch, K., Klapproth, K., Barile, M., Flossdorf, M., Holland-Letz, T., Schlenner, S. M., Reth, M., Höfer, T. and Rodewald, H. R. (2015). Fundamental properties of unperturbed haematopoiesis from stem cells in vivo. Nature 518, 542–546.

Cabezas-Wallscheid, N., Buettner, F., Sommerkamp, P., Klimmeck, D., Ladel, L., Thalheimer, F. B., Pastor-Flores, D., Roma, L. P., Renders, S., Zeisberger, P., Przybylla, A., Schonberger, K., Scognamiglio, R., Altamura, S., Florian, C. M., Fawaz, M., Vonficht, D., Tesio, M., Collier, P., Pavlinic, D., Geiger, H., Schroeder, T., Benes, V., Dick, T. P., Rieger, M. A., Stegle, O. and Trumpp, A. (2017). Vitamin A-Retinoic Acid Signaling Regulates Hematopoietic Stem Cell Dormancy. Cell 169, 807–823.

Carrelha, J., Meng, Y., Kettyle, L. M., Luis, T. C., Norfo, R., Alcolea, V., Boukarabila, H., Grasso, F., Gambardella, A., Grover, A., Hogstrand, K., Lord, A. M., Sanjuan-Pla, A., Woll, P. S., Nerlov, C. and Jacobsen, S. E. W. (2018). Hierarchically related lineage-restricted fates of multipotent haematopoietic stem cells. Nature 554, 106–111.

Chapple, R. H., Tseng, Y. J., Hu, T., Kitano, A., Takeichi, M., Hoegenauer, K. A. and Nakada, D. (2018). Lineage tracing of murine adult hematopoietic stem cells reveals active contribution to steady-state hematopoiesis. Blood Adv 2, 1220–1228.

De Boer, R. J. and Perelson, A. S. (2013). Quantifying T lymphocyte turnover. J. Theor. Biol. 327, 45–87.

Domen, J., Cheshier, S. H. and Weissman, I. L. (2000). The role of apoptosis in the regulation of hematopoietic stem cells: Overexpression of Bcl-2 increases both their number and repopulation potential. J. Exp. Med. 191, 253–264.

Florian, M. C., Klose, M., Sacma, M., Jablanovic, J., Knudson, L., Nattamai, K. J., Marka, G., Vollmer, A., Soller, K., Sakk, V., Cabezas-Wallscheid, N., Zheng, Y., Mulaw, M. A., Glauche, I. and Geiger, H. (2018). Aging alters the epigenetic asymmetry of HSC division. PLoS Biol. 16, e2003389.

Foudi, A., Hochedlinger, K., Van Buren, D., Schindler, J. W., Jaenisch, R., Carey, V. and Hock, H. (2009). Analysis of histone 2B-GFP retention reveals slowly cycling hematopoietic stem cells. Nat. Biotechnol. 27, 84–90.

Gao, J., Van Meter, M., Hernandez Lopez, S., Chen, G., Huang, Y., Ren, S., Zhao, Q., Rojas, J., Gurer, C., Thurston, G. and Kuhnert, F. (2019). Therapeutic targeting of Notch signaling and immune checkpoint blockade in a spontaneous, genetically heterogeneous mouse model of T-cell acute lymphoblastic leukemia. Dis Model Mech 12.

Gazit, R., Mandal, P. K., Ebina, W., Ben-Zvi, A., Nombela-Arrieta, C., Silberstein, L. E. and Rossi, D. J. (2014). Fgd5 identifies hematopoietic stem cells in the murine bone marrow. J. Exp. Med. 211, 1315–1331.

Giladi, A., Paul, F., Herzog, Y., Lubling, Y., Weiner, A., Yofe, I., Jaitin, D., Cabezas-Wallscheid, N., Dress, R., Ginhoux, F., Trumpp, A., Tanay, A. and Amit, I. (2018). Single-cell characterization of haematopoietic progenitors and their trajectories in homeostasis and perturbed haematopoiesis. Nat. Cell Biol. 20, 836–846.

Haghverdi, L., Buttner, M., Wolf, F. A., Buettner, F. and Theis, F. J. (2016). Diffusion pseudo-time robustly reconstructs lineage branching. Nat. Methods 13, 845–848.

Höfer, T. and Rodewald, H. R. (2016). Output without input: the lifelong productivity of hematopoietic stem cells. Curr. Opin. Cell Biol. 43, 69–77.

Inaba, M. and Yamashita, Y. M. (2012). Asymmetric stem cell division: precision for robustness. Cell Stem Cell 11, 461–469.

Ito, K., Turcotte, R., Cui, J., Zimmerman, S. E., Pinho, S., Mizoguchi, T., Arai, F., Runnels, J. M., Alt, C., Teruya-Feldstein, J., Mar, J. C., Singh, R., Suda, T., Lin, C. P., Frenette, P. S. and Ito, K. (2016). Self-renewal of a purified Tie2+ hematopoietic stem cell population relies on mitochondrial clearance. Science 354, 1156–1160.

Karamitros, D., Stoilova, B., Aboukhalil, Z., Hamey, F., Reinisch, A., Samitsch, M., Quek, L., Otto, G., Repapi, E., Doondeea, J., Usukhbayar, B., Calvo, J., Taylor, S., Goardon, N., Six, E., Pflumio, F., Porcher, C., Majeti, R., G?ttgens, B. and Vyas, P. (2018). Single-cell analysis reveals the continuum of human lympho-myeloid progenitor cells. Nat. Immunol. 19, 85–97.

Kiel, M. J., He, S., Ashkenazi, R., Gentry, S. N., Teta, M., Kushner, J. A., Jackson, T. L. and Morrison, S. J. (2007). Haematopoietic stem cells do not asymmetrically segregate chromosomes or retain BrdU. Nature 449, 238–242.

Klein, A. M. and Simons, B. D. (2011). Universal patterns of stem cell fate in cycling adult tissues. Development 138, 3103–3111.

Lajtha, L. G., Olivier, R. and Gurney, C. W. (1962). Kinetic model of a bone-marrow stem-cell population. Br. J. Haematol. 8, 442–460.

Laurenti, E. and Gottgens, B. (2018). From haematopoietic stem cells to complex differentiation landscapes. Nature 553, 418–426.

Loeffler, D., Wehling, A., Schneiter, F., Zhang, Y., Muller-Botticher, N., Hoppe, P. S., Hilsenbeck, O., Kokkaliaris, K. D., Endele, M. and Schroeder, T. (2019). Asymmetric lysosome inheritance predicts activation of haematopoietic stem cells. Nature 573, 426–429.

Mackey, M. C. (2001). Cell kinetic status of haematopoietic stem cells. Cell Prolif. 34, 71–83.

Martins, V. C., Busch, K., Juraeva, D., Blum, C., Ludwig, C., Rasche, V., Lasitschka, F., Mastitsky, S. E., Brors, B., Hielscher, T., Fehling, H. J. and Rodewald, H. R. (2014). Cell competition is a tumour suppressor mechanism in the thymus. Nature 509, 465–470.

Mende, N., Jolly, A., Percin, G. I., Günther, M., Rostovskaya, M., Krishnan, S. M., Oostendorp, R. A. J., Dahl, A., Anastassiadis, K., Höfer, T. and Waskow, C. (2019). Prospective isolation of nonhematopoietic cells of the niche and their differential molecular interactions with HSCs. Blood 134, 1214–1226.

Morcos, M. N. F., Zerjatke, T., Glauche, I., Munz, C. M., Ge, Y., Petzold, A., Reinhardt, S., Dahl, A., Anstee, N. S., Bogeska, R., Milsom, M. D., S?w?n, P., Wan, H., Bryder, D., Roers, A. and Gerbaulet, A. (2020). Continuous mitotic activity of primitive hematopoietic stem cells in adult mice. J. Exp. Med. 217.

Morrison, S. J. and Kimble, J. (2006). Asymmetric and symmetric stem-cell divisions in development and cancer. Nature 441, 1068–1074.

Morrison, S. J. and Spradling, A. C. (2008). Stem cells and niches: mechanisms that promote stem cell maintenance throughout life. Cell 132, 598–611.

Morrison, S. J., Wandycz, A. M., Akashi, K., Globerson, A. and Weissman, I. L. (1996). The aging of hematopoietic stem cells. Nat. Med. 2, 1011–1016.

Nestorowa, S., Hamey, F. K., Pijuan Sala, B., Diamanti, E., Shepherd, M., Laurenti, E., Wilson, N. K., Kent, D. G. and G?ttgens, B. (2016). A single-cell resolution map of mouse hematopoietic stem and progenitor cell differentiation. Blood 128, 20–31.

Oguro, H., Ding, L. and Morrison, S. J. (2013). SLAM family markers resolve functionally distinct subpopulations of hematopoietic stem cells and multipotent progenitors. Cell Stem Cell 13, 102–116.

Olsson, A., Venkatasubramanian, M., Chaudhri, V. K., Aronow, B. J., Salomonis, N., Singh, H. and Grimes, H. L. (2016). Single-cell analysis of mixed-lineage states leading to a binary cell fate choice. Nature 537, 698–702.

Osgood, E. E. (1957). A unifying concept of the etiology of the leukemias, lymphomas, and cancers. J. Natl. Cancer Inst. 18, 155–166.

Paul, F., Arkin, Y., Giladi, A., Jaitin, D. A., Kenigsberg, E., Keren-Shaul, H., Winter, D., Lara-Astiaso, D., Gury, M., Weiner, A., David, E., Cohen, N., Lauridsen, F. K., Haas, S., Schlitzer, A., Mildner, A., Ginhoux, F., Jung, S., Trumpp, A., Porse, B. T., Tanay, A. and Amit, I. (2015). Transcriptional Heterogeneity and Lineage Commitment in Myeloid Progenitors. Cell 163, 1663–1677.

Pei, W., Feyerabend, T. B., Rossler, J., Wang, X., Postrach, D., Busch, K., Rode, I., Klapproth, K., Dietlein, N., Quedenau, C., Chen, W., Sauer, S., Wolf, S., Höfer, T. and Rodewald, H. R. (2017). Polylox barcoding reveals haematopoietic stem cell fates realized in vivo. Nature 548, 456–460.

Pei, W., Shang, F., Wang, X., Fanti, A.-K., Busch, K., Klapproth, K., Zhang, Q., Quedenau, C., Sauer, S., Feyerabend, T. B., Höfer, T. and Rodewald, H.-R. (2020). Resolving fate and transcriptome of hematopoietic stem cell clones. BioRxiv 2020.03.25.008433.

Rodriguez-Fraticelli, A. E., Wolock, S. L., Weinreb, C. S., Panero, R., Patel, S. H., Jankovic, M., Sun, J., Calogero, R. A., Klein, A. M. and Camargo, F. D. (2018). Clonal analysis of lineage fate in native haematopoiesis. Nature 553, 212–216.

Rulands, S. and Simons, B. D. (2016). Tracing cellular dynamics in tissue development, maintenance and disease. Curr. Opin. Cell Biol. 43, 38–45.

Sawai, C. M., Babovic, S., Upadhaya, S., Knapp, D. J. H. F., Lavin, Y., Lau, C. M., Goloborodko, A., Feng, J., Fujisaki, J., Ding, L., Mirny, L. A., Merad, M., Eaves, C. J. and Reizis, B. (2016). Hematopoietic Stem Cells Are the Major Source of Multilineage Hematopoiesis in Adult Animals. Immunity 45, 597–609.

Säwen, P., Eldeeb, M., Erlandsson, E., Kristiansen, T. A., Laterza, C., Kokaia, Z., Karlsson, G., Yuan, J., Soneji, S., Mandal, P. K., Rossi, D. J. and Bryder, D. (2018). Murine HSCs contribute actively to native hematopoiesis but with reduced differentiation capacity upon aging. Elife 7.

Schoedel, K. B., Morcos, M. N. F., Zerjatke, T., Roeder, I., Grinenko, T., Voehringer, D., Gothert, J. R., Waskow, C., Roers, A. and Gerbaulet, A. (2016). The bulk of the hematopoietic stem cell population is dispensable for murine steady-state and stress hematopoiesis. Blood 128, 2285–2296.

Sheikh, B. N., Yang, Y., Schreuder, J., Nilsson, S. K., Bilardi, R., Carotta, S., McRae, H. M., Metcalf, D., Voss, A. K. and Thomas, T. (2016). MOZ (KAT6A) is essential for the maintenance of classically defined adult hematopoietic stem cells. Blood 128, 2307–2318.

Simons, B. D. and Clevers, H. (2011). Strategies for homeostatic stem cell self-renewal in adult tissues. Cell 145, 851–862.

Sudo, K., Ema, H., Morita, Y. and Nakauchi, H. (2000). Age-associated characteristics of murine hematopoietic stem cells. J. Exp. Med. 192, 1273–1280.

Sun, J., Ramos, A., Chapman, B., Johnnidis, J. B., Le, L., Ho, Y. J., Klein, A., Hofmann, O. and Camargo, F. D. (2014). Clonal dynamics of native haematopoiesis. Nature 514, 322–327.

van der Wath, R. C., Wilson, A., Laurenti, E., Trumpp, A. and Lio, P. (2009). Estimating dormant and active hematopoietic stem cell kinetics through extensive modeling of bromodeoxyuridine label-retaining cell dynamics. PLoS ONE 4, e6972.

Velten, L., Haas, S. F., Raffel, S., Blaszkiewicz, S., Islam, S., Hennig, B. P., Hirche, C., Lutz, C., Buss, E. C., Nowak, D., Boch, T., Hofmann, W. K., Ho, A. D., Huber, W., Trumpp, A., Essers, M. A. and Steinmetz, L. M. (2017). Human haematopoietic stem cell lineage commitment is a continuous process. Nat. Cell Biol. 19, 271–281.

Wilson, A., Laurenti, E., Oser, G., van der Wath, R. C., Blanco-Bose, W., Jaworski, M., Offner, S., Dunant, C. F., Eshkind, L., Bockamp, E., Lio, P., Macdonald, H. R. and Trumpp, A. (2008). Hematopoietic stem cells reversibly switch from dormancy to self-renewal during homeostasis and repair. Cell 135, 1118–1129.

